# Whole genome sequencing in multiplex families reveals novel inherited and *de novo* genetic risk in autism

**DOI:** 10.1101/338855

**Authors:** Elizabeth K. Ruzzo, Laura Pérez-Cano, Jae-Yoon Jung, Lee-kai Wang, Dorna Kashef-Haghighi, Chris Hartl, Jackson Hoekstra, Olivia Leventhal, Michael J. Gandal, Kelley Paskov, Nate Stockham, Damon Polioudakis, Jennifer K. Lowe, Daniel H. Geschwind, Dennis P. Wall

## Abstract

Genetic studies of autism spectrum disorder (ASD) have revealed a complex, heterogeneous architecture, in which the contribution of rare inherited variation remains relatively un-explored. We performed whole-genome sequencing (WGS) in 2,308 individuals from families containing multiple affected children, including analysis of single nucleotide variants (SNV) and structural variants (SV). We identified 16 new ASD-risk genes, including many supported by inherited variation, and provide statistical support for 69 genes in total, including previously implicated genes. These risk genes are enriched in pathways involving negative regulation of synaptic transmission and organelle organization. We identify a significant protein-protein interaction (PPI) network seeded by inherited, predicted damaging variants disrupting highly constrained genes, including members of the BAF complex and established ASD risk genes. Analysis of WGS also identified SVs effecting non-coding regulatory regions in developing human brain, implicating *NR3C2* and a recurrent 2.5Kb deletion within the promoter of *DLG2*. These data lend support to studying multiplex families for identifying inherited risk for ASD. We provide these data through the Hartwell Autism Research and Technology Initiative (iHART), an open access cloud-computing repository for ASD genetics research.

## Main text

Autism spectrum disorder (ASD) is a neurodevelopmental disorder characterized by early deficits in social communication and interaction, together with restricted and repetitive patterns of behavior, interest, or activity(*1*). Global prevalence is between 1-2%(*2*), and ASD has a strong genetic component, with heritability estimated between 60% to 90%(*3–9*). Considerable progress in gene discovery has come from studies in families with only one affected child (simplex families) identifying *de novo* (DN) copy number variants (CNV)(*10-13*) and *de novo* frameshift, splice-acceptor, splice-donor, or nonsense variants (collectively referred to as protein-truncating variants (PTVs))(*14–18*) that increase ASD-risk and account for an estimated 3-5% of ASD cases(*7, 8, 19–21*). Despite having identified roughly 90 ASD-risk genes with high confidence(*22, 23*), it is estimated that between 260 and 1,250 genes confer risk for developing ASD(*24*), leaving substantial room for gene discovery, especially with regards to inherited variation. To date, several recurrent CNVs have also been associated with increased ASD risk(*25–27*). Evidence for inherited risk variants has been drawn primarily from families containing only one affected child(*18, 28*), which are depleted for inherited risk as compared to families with two or more affected children (multiplex families)(*10, 24, 29*). Here we used WGS to study the coding and non-coding regions of the genome in the largest cohort of multiplex families evaluated to date to identify *de novo* and inherited genetic risk factors for ASD.

## Results

WGS was performed to an average depth of 35.9 ± 5.0 (**Fig. S1; Methods**) in 2,308 individuals (n=493 families) from the Autism Genetic Resource Exchange (AGRE); purified DNA was sequenced from whole blood (n=285) or lymphoblastoid cell line (LCL) DNA (n=2,023) when whole blood DNA was not available. Our cohort consists of families with two or more children diagnosed with ASD and excludes families with known genetic causes or syndromes (**Methods; Table S1, Table S2**). Single nucleotide variants (SNV) and indels were identified following GATK’s best practices(*30*) (**Methods**) and structural variants (SV) were identified using a combination of BreakDancer(*31*), GenomeSTRiP(*32, 33*), LUMPY(*34*), and Somatic MUtation FINder (SMuFin)(*35*) (**Methods; Fig. 1, Fig. S2**). We adapted SMuFin for family-based structural variant detection by performing *de novo* alignment of child reads to the parental reads (**Fig. S2c**) to provide high sensitivity and break point accuracy in the detection of SVs (**Methods**). After quality control (**Methods**), we categorized the inheritance of each variant in the children (e.g., *de novo*, paternally inherited, unknown phase) and annotated variant/gene properties to facilitate downstream analyses (**Fig. 1**).

**Fig. 1.**
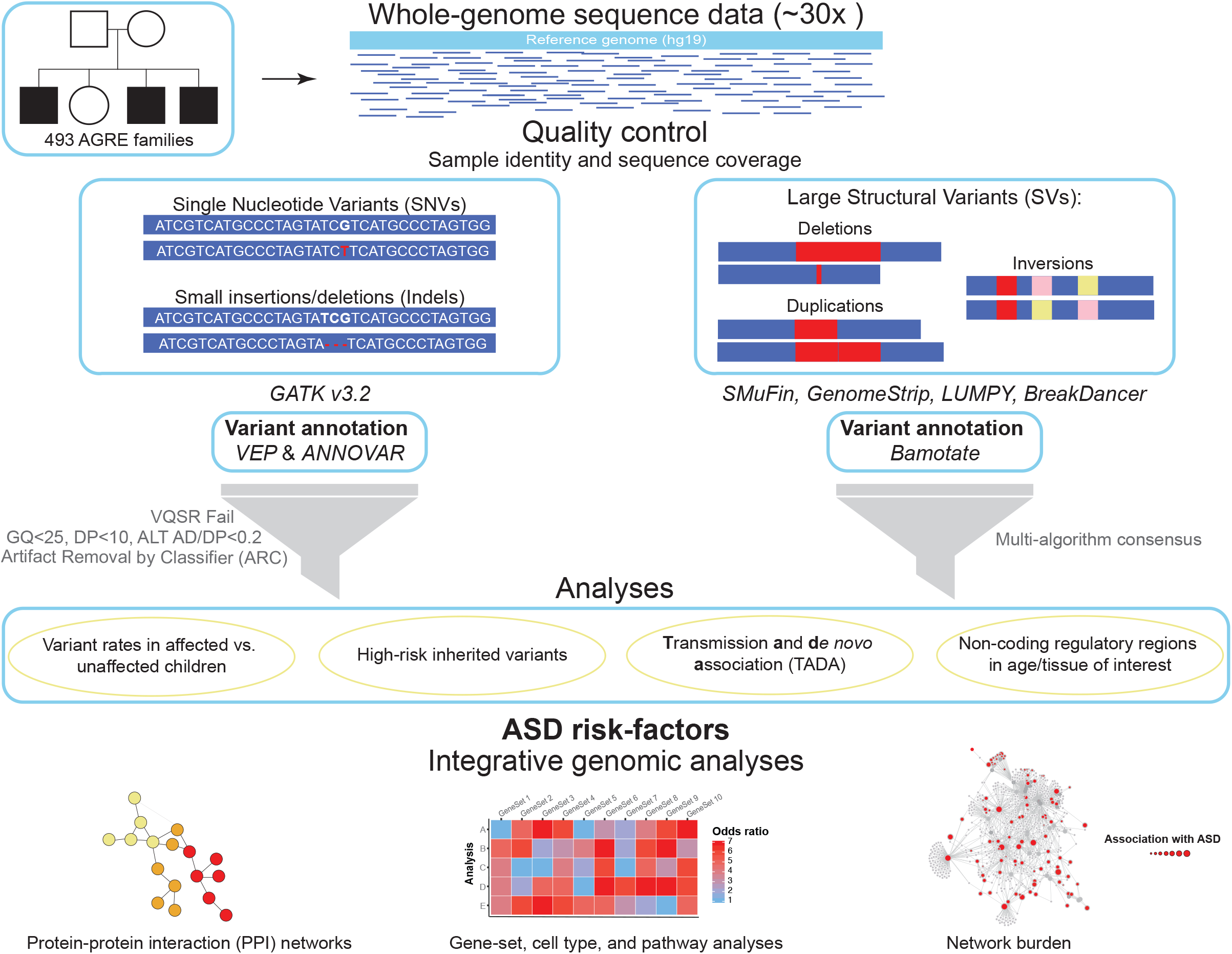
Overview of the Analysis Pipeline. High-coverage (30x) whole-genome sequence data was generated at the NYGC on the Illumina HiSeq X. Reads were aligned to the human reference genome (hg19) and single nucleotide variants and indels were called following GATK’s best practices. Quality control checks were applied to insure both sequencing/variant quality and sample identity (Methods, Fig. S1). SNVs and indels were annotated using both VEP(*71*) and ANNOVAR(*70*) and subsequently filtered for mildly stringent quality thresholds. All *de novo* variants were subjected to our machine learning classifier – Artifact Removal by Classifier (ARC; Methods, Fig. 3, Fig. S5-S8). Large structural variants were identified by four different SV-detection algorithms (Methods) – three of which used aligned sequence reads and one that performed *de novo* alignment (SMuFin). Large SVs were annotated using Bamotate and then filtered for high quality variants by using our multi-algorithm consensus pipeline (Methods). The resulting variants were then analyzed to identify ASD-risk factors and perform integrative genomic analyses.

### Inherited coding and promoter-disrupting variants highlight a syndromic form of ASD and a novel deletion associated with ASD

Since multiplex ASD families are expected to be enriched for inherited risk variants(*10, 24, 29*), we first assessed rare (allele frequency (AF) ≤0.1%) inherited variants, finding no excess of rare inherited missense or PTV variants in affected subjects (**Fig. 2a, Fig. S3**). Based on previous analyses(*28*), we asked if there was an enrichment of inherited private PTVs in loss-of-function intolerant genes (pLI≥0.9)(*36*) and found no significant excess in affected subjects (*P*=0.40, quasi-Poisson linear regression, **Fig. S4a**). We also observed no difference in the overall rate of rare inherited SVs between affected and unaffected individuals, even when restricting to gene-disrupting SVs (**Fig. S4c-h**).

**Fig. 2.**
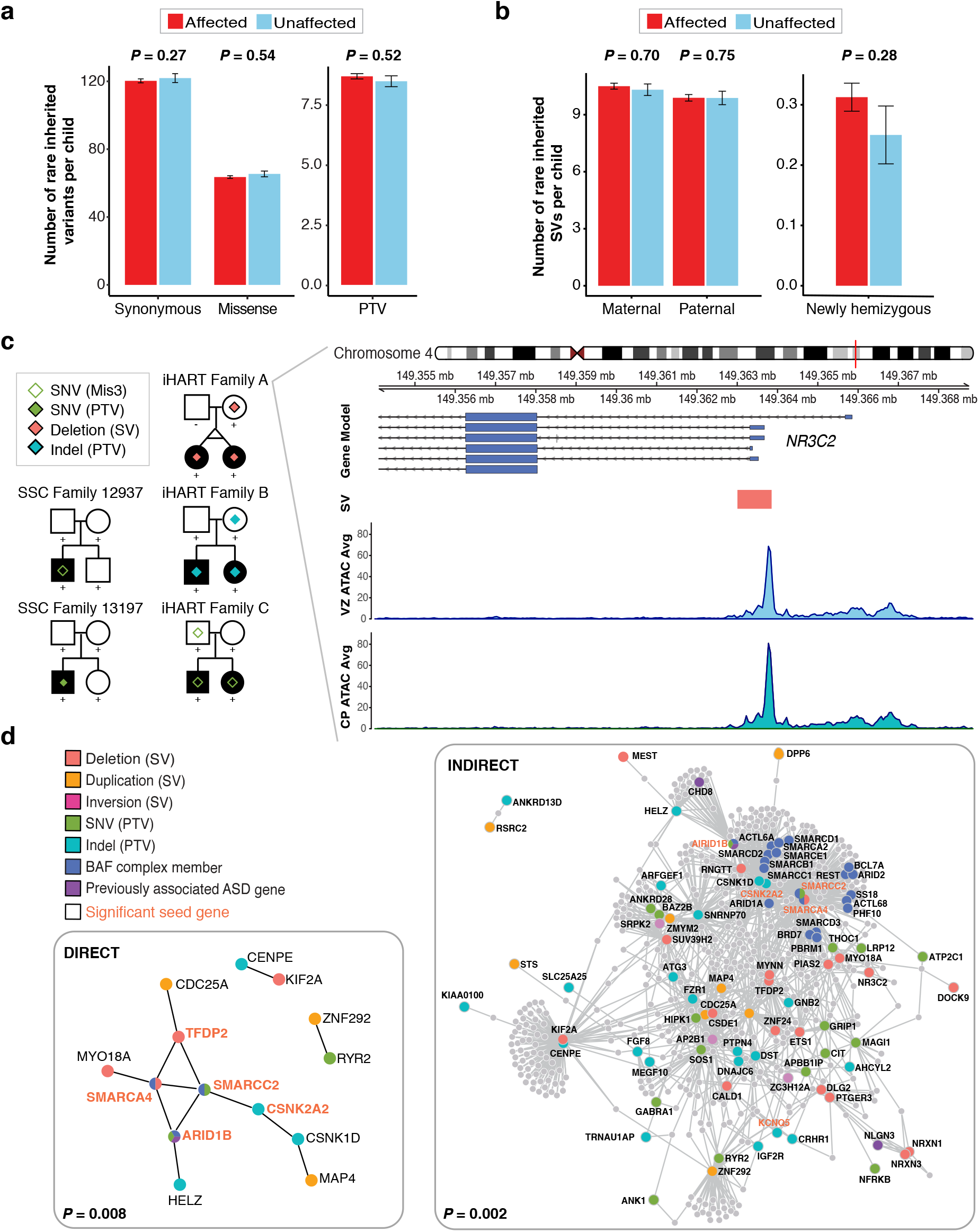
Inherited ASD-risk genes. **(a)** The rate of rare inherited coding variants per fully phase-able child is displayed for 960 affected (red) and 217 unaffected children (light blue) by variant consequence. Rare inherited variants (SNVs and indels) were defined as those with an AF≤0.1% in public databases (1000g, ESP6500, ExACv3.0, cg46), internal controls, and iHART HNP samples and were restricted to those not missing in more than 25% of controls and that were not flagged as low-confidence by the GIAB consortium (Methods). Rates are displayed as bar plots of the mean number of variants in a sample and error bars represent the standard error. **(b)** The rate of rare inherited SVs per fully phase-able child is displayed for 960 affected (red) and 217 unaffected children (light blue) by inheritance type; this includes newly hemizygous variants in 563 affected (red) and 100 unaffected male children (light blue). Rare SVs (DELs, DUPs, INVs) were defined as those with an AF < 0.001 in cDGV and an AF < 0.01 in iHART HNP samples. **(c)** Pedigrees are displayed for five ASD families identified as harboring coding or regulatory variants for *NR3C2*. Both SSC families harbor *de novo* variants in the proband which are absent in the unaffected sibling (a probably damaging missense (Mis3) in SSC12937 and a PTV in SSC13197). AGRE/iHART families A-C harbor rare inherited variants transmitted to both affected children (a promoter-disrupting deletion in family A, a PTV in family B, and a probably damaging missense (Mis3) in family C). The *NR3C2* promoter-disrupting ~850bp deletion (orange rectangle) transmitted to all affected members of family A. Shown below are the average ATAC-seq peak read depth from the cortical plate (CP) and ventricular zone (VZ) of developing human brain samples (n=3); **(d)** The direct and indirect protein-protein interaction networks (identified by DAPPLE) formed by intolerant genes harboring PTVs or damaging SVs (promoter or exon disrupting) transmitted to all affected (and no unaffected) members of an AGRE family. Each encoded protein is colored according to the type of variant class identified. Proteins encoded by an ASD risk gene identified in the TADA mega analysis in Sanders *et al*. (*22*) are shown in purple. Any protein falling in more than one category is colored with all categorical colors that apply. The gene labels for significant seed genes are bold and orange.

We next identified rare, damaging variants in loss-of-function intolerant genes (pLI≥0.9)(*36*) transmitted to all affected individuals of a multiplex family, but not to the unaffected individuals (**Methods**). A total of 62 PTVs and 40 SVs disrupting a coding exon or promoter were identified in 98 unique genes; including three genes, *NR3C2, NRXN1*, and *ZMYM2*, disrupted by both a PTV in one family and an SV in a second family. For example, the *NR3C2* gene harbors a rare PTV and a ~850bp deletion (chr4: 149363005 - 149363852) in the promoter region of *NR3C2* that are transmitted to both affected children in each family (**Fig. 2c**). The deleted promoter region of *NR3C2* falls in a functional non-coding regulatory region in developing human brain(*37*) (chr4:149362706-149367485) (**Fig. 2c**). Expanding to include PolyPhen-2 damaging missense variants in *NR3C2*, we identify a third family with a rare missense variant transmitted from father to both affected children (**Fig. 2c**). The three families identified with a transmitted regulatory or protein-disrupting variant in *NR3C2* share striking phenotypic similarities, defining a new syndromic form of ASD characterized by metacarpal hypoplasia, high arched palate, sensory hypersensitivity, and abnormal prosody (**Table S3**).

Additionally, three families carry the same 2.5Kb deletion in the promoter region of *DLG2* (chr11: 85339733 – 85342186); a gene that is associated with cognition and learning in mice and humans(*38, 39*) (**Fig. S4b**). This deletion falls in a previously-defined functional non-coding regulatory region in developing human brain(*37*) (chr11: 85338026 - 85340560) (**Fig. S4b**) and occurs on a different haplotype in each of the three families (**Methods, Table S4**). No deletions were observed to overlap the identified *DLG2* promoter deletion in public databases or any controls (n = 26,353 controls, **Methods**). This rare regulatory mutation is significantly associated with ASD (3 of 484 unrelated affected children versus 0 of 26,565 controls, two-sided Fisher’s Exact Test, *P*=5.7 × 10^−6^, OR=Inf, 95% CI=22.7-Inf, **Methods**); this association remains significant even when considering only WGS control samples (**Methods**, n=2,889 controls, twosided Fisher’s Exact Test, *P*=0.003, OR=Inf, 95% CI=2.47-Inf).

### No association signal is observed in promoters of established ASD risk genes

To further assess the role of rare regulatory variation in ASD in addition to SVs, we also assessed the impact of non-coding SNVs or indels in promoters (**Methods**), observing no enrichment for RDNVs in affected vs. unaffected iHART children when looking globally (**Methods**; *P*=0.33, quasi-Poisson linear regression), or when restricting the analysis to known ASD-risk genes (**Methods**; *P*=0.42, quasi-Poisson linear regression). Similarly, we see no signal for private inherited variants in promoters (**Methods**; All genes *P*=0.07 and ASD-risk genes *P*=0.26, quasi-Poisson linear regression), nor when our cohort is combined with 517 affected and 518 unaffected children with WGS data from the Simons Simplex Cohort (SSC) (**Methods**; RDNVs: All genes *P*=0.25 and ASD-risk genes *P*=0.31, quasi-Poisson linear regression; Private inherited: All genes *P*=0.14 and ASD-risk genes *P*=0.12). These data are consistent with recent results in simplex families(*40*).

### Genes hit by high-risk inherited variants show biological convergence

We next determined whether the 98 genes harboring PTVs and SVs transmitted to all affected individuals represented a random or broadly acting collection of genes, or had evidence for biological convergence, reasoning that the latter would provide orthogonal support. Indeed, the protein products of these genes formed a significant direct protein–protein interaction (PPI) network (**Methods**, 1000 permutations, *P* < 0.008; **Fig. 2d**). This network is enriched for members of the BAF (SWI/SNF) complex (two-sided Fisher’s Exact Test, *P*=0.02, OR=5.9, 95% confidence interval 1.1 – 20.7), including *ARID1B, SMARCC2*, and *SMARCA4*, which are involved in chromatin remodeling during cortical neurogenesis and have previously been associated with ASD(*41, 42*). Given that PPI databases are incomplete and biased against typically less well-studied neuronal interactions(*43*), we expanded the network to include indirect interactions among the seed genes. This indirect interaction network was also significant (*P* < 0.002) and seven proteins in this network were found to be significantly connected hubs (corrected seed score *P* < 0.05) (**Fig. 2d**). Gene set enrichment (**Methods**) identified slight enrichment for targets of RBFOX1(*44*) (*P*=0.034, uncorrected), which regulates neuronal alternative splicing and previously has been implicated in ASD(*10, 45*).

### Identification of high-quality *de novo* variants by machine learning

*De novo* missense and PTVs have been identified as significant risk factors for ASD in simplex families(*17, 18, 46*). To investigate the role of these variant classes in patients with ASD from multiplex families, we developed a supervised random forest model, Artifact Removal by Classifier (ARC), to distinguish true rare *de novo* variants (RDNVs) from LCL-specific genetic aberrations or other types of artifacts, such as sequencing and mapping errors. We trained ARC on RDNVs identified in 76 pairs of fully phase-able monozygotic (MZ) twins with WGS data derived from LCL DNA, using 48 features representing intrinsic genomic properties, (e.g., GC content and properties associated with *de novo* hotspots(*47*)), sample specific properties (e.g., genome-wide number of *de novo* SNVs), signatures of transformation of peripheral B lymphocytes by Epstein-Barr virus (e.g., number of *de novo* SNVs in immunoglobulin genes), or variant properties (e.g., GATK variant metrics) (**Fig. 3c; Table S5**). We subsequently tested ARC on RDNVs identified in 17 fully phase-able whole blood (WB) and matched LCL samples with WGS data (**Fig. S5**). The resulting random forest classifier achieved an area under the receiver operating characteristic (ROC) curve of 0.99 and 0.98 in the training and test set, respectively (**Fig. 3a, 3b**). We selected a conservative ARC score threshold (0.4) that achieved a minimum precision and recall rate of >0.9 and ~0.8, respectively, across all 10-folds of the training set cross validation (**Fig. S7c, S7d**); and achieved a precision and recall rate of >0.9 and >0.8, respectively, in the test set (**Fig. S7h**).

**Fig. 3.**
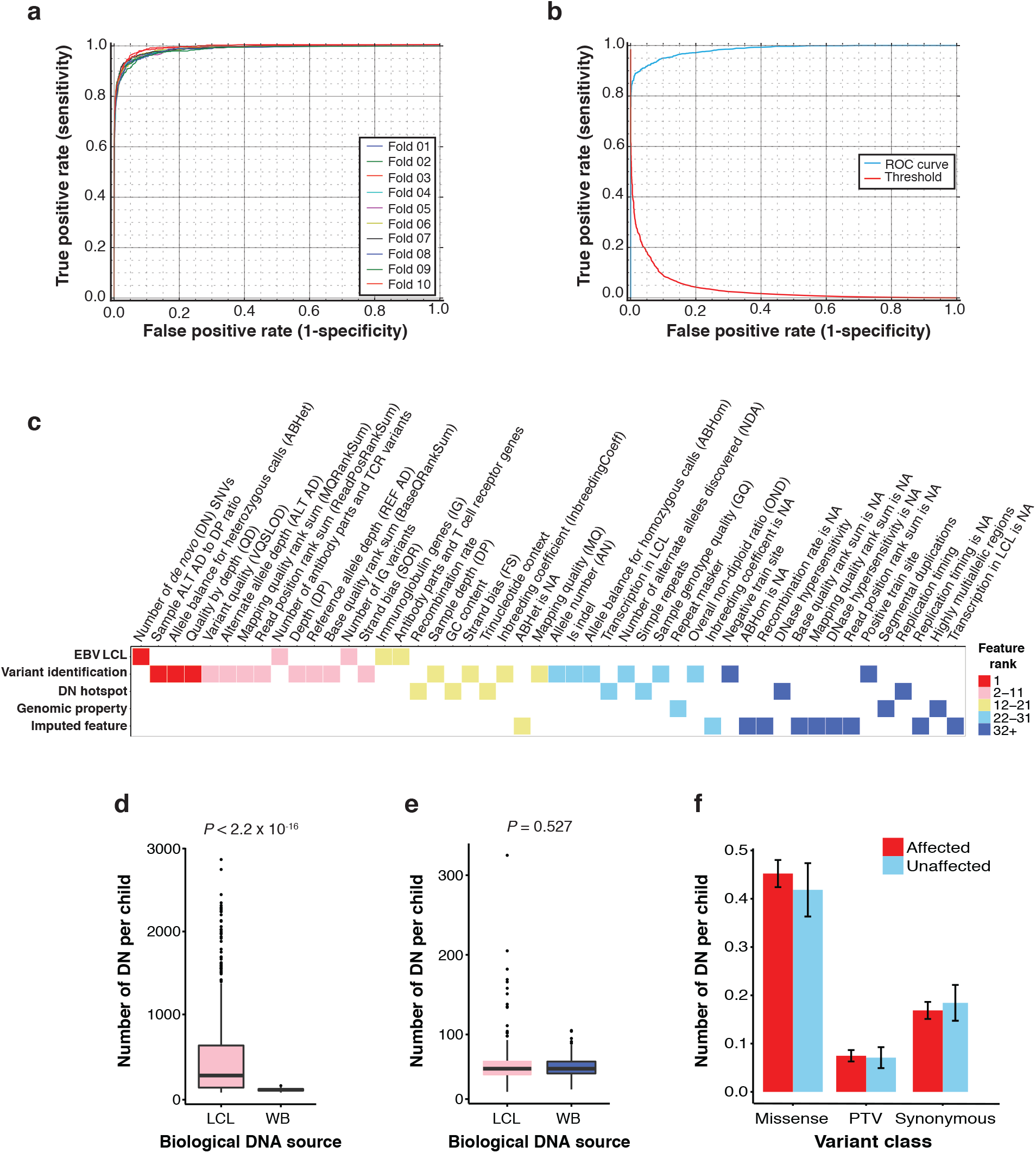
Rare *de novo* variants in iHART/AGRE. **(a)** The ROC curves for the 10-fold cross validation for the ARC training set; AUC=0.99. **(b)** The ROC curve for the ARC test set; AUC=0.98. **(c)** The relative importance of each random forest feature was obtained from the RFECV module from scikit-learn. This heat map reflects the rank of all 48 ARC features (binned for heat map coloring but listed on the x-axis in order of rank) and displays features by relevant categories (y-axis). **(d)** The number of rare *de novo* variants identified in LCL (pink) and WB (blue) fully phase-able (non-MZ twin) samples before ARC (N=1,019 samples). The difference in DN rates between the biological sequencing source (LCL vs. WB) was evaluated using Wilcoxon rank sum test. **(e)** The number of rare *de novo* variants identified in LCL (pink) and WB (blue) fully phase-able (non-MZ twin) samples after ARC (variants with an ARC score <0.4 are filtered out) and after excluding ARC outlier samples (samples with >90% DNs removed by ARC) (N=716). No significant difference in the rate of rare *de novo* variants based on the biological sequencing source (LCL_median_=57 and WB_median_=57) using the Wilcoxon rank sum test. **(f)** The rate of rare *de novo* coding variants per child is displayed for 575 affected (red) and 141 unaffected children (light blue) (716 fully phase-able samples after excluding MZ twins and ARC outliers) by variant consequence. Rare *de novo* variants (absent in all controls) were restricted to those with an ARC score ≥0.4 that were not flagged as low-confidence by the GIAB consortium (Methods). Rates are displayed as bar plots of the mean number of rare *de novo* variants in a sample and error bars represent the standard error.

We applied ARC to all raw RDNVs identified in the 1,177 children for whom both biological parents were also sequenced (fully phase-able samples) from 422 iHART families, observing that ARC eliminated many more raw RDNVs in LCL samples than in WB samples (**Fig. S8a**) and resulted in the elimination of a significant difference in RDNV rates between biological sequencing sources (before ARC: FET *P* = <2.2×10^−16^; after ARC: FET *P* = 0.527; **Fig. 3d, 3e** and **Fig. S8b**). Use of ARC yielded a mean genome-wide *de novo* mutation rate of 60.3 RDNVs per child in LCL-derived samples and 59.4 RDNVs per child in WB-derived samples (**Fig. 3e**), on the lower end of reported genome-wide *de novo* mutation rates (mean=64.4; range 54.8-81)(*23, 47–51*), consistent with our conservative approach. Further support for ARC’s performance comes from the observation that the effect of paternal age on the number of *de novo* mutations per affected ASD child is seen only after running ARC (*P*=3.6 × 10^−13^). After application of ARC our observed rate of 1.46 RDNVs per year of paternal age (95% CI = 1.37-1.55) matches previously published rates(*47, 52–54*) (**Methods; Fig. S8c**).

### Evidence for depletion of rare *de novo* ASD risk variants in multiplex families

Neither *de novo* missense (*P*=0.561, quasi-Poisson linear regression) nor PTVs (*P*=0.873, quasi-Poisson linear regression) showed a significant association in iHART/AGRE multiplex families (**Fig. 3f**). Our finding is consistent with a previous well-powered study of rare *de novo* copy number variants (CNVs) which showed significant enrichment in affected children as compared to unaffected siblings in simplex families from the SSC, but not in multiplex families from AGRE(*27*). The rate of rare *de novo* PTVs in affected children from multiplex families in our study (Aff_iHART_=0.07) was approximately half of that in simplex families (Aff_Kosmicki_=0.13)(17, 55) (Table S6). Furthermore, the rate of rare *de novo* PTVs in unaffected children from multiplex families (Unaff_iHART_=0.07) was identical to that of affected children; by Monte Carlo integration, we estimated that our current cohort had >70% power to detect a rate difference for *de novo* PTVs in affected and unaffected children (**Methods**). In contrast, comparable rates of rare *de novo* synonymous and missense variants in both affected and unaffected children were observed regardless of family structure (**Table S6**).

### Identification of 16 novel ASD risk genes enriched for inherited variation

We next used the Transmitted And *De novo* Association (TADA) test(*56*) to increase power for gene discovery by combining evidence from rare *de novo* (DN) or transmitted (inherited) protein-truncating variants (PTVs) and *de novo* missense variants predicted to damage the encoded protein (Mis3, a probably damaging prediction by PolyPhen-2(*57*)) (**Methods**). Since the distribution of the TADA statistic (under the null) is not known for multiplex families, we estimated the distribution of the null TADA statistic by simulating Mendelian transmission and *de novo* mutation across family structures (**Methods**). To further improve power, we combined qualifying variants found in ASD samples from the current (iHART) cohort (**Table S7**) with the most recent ASD TADA mega-analysis(*22*) (**Table S8**), identifying 69 genes reaching the threshold of FDR<0.1, (**Table 1, Fig. 4a, Table S9-S11**), of which 16 had not previously been identified as ASD-risk genes (**Table 1, Fig. 4a**). These 16 new risk genes are enriched for those in which a higher proportion of risk variants are inherited (**Methods**). For six of the 16 novel genes (*UIMC1, C16orf13, MLANA, CCSER1, PCM1, FAM98C*) and five of the 53 previously associated (*RANBP17, ZNF559, P2RX5, CTTNBP2, CAPN12*) ASD-risk genes, ≥70% of the qualifying variants are inherited PTV (Fisher’s Exact Test, *P* = 0.015, OR=5.57, 95% CI:1.17-28.35). Remarkably, for *PCM1*, the Bayes Factor contribution from inherited PTVs was greater than the Bayes Factor for *de novo* variants (**Methods**), indicating that the association signal is mainly driven by inherited PTVs. To ensure that we were not obtaining type I errors due to family structure alone, we also performed simulations of the distribution of the null TADA statistic using the observed variant counts (**Methods**). Genes with the lowest FDR in the TADA-mega analysis showed the largest simulated Bayes factors and lowest p-values (**Fig. S9**). All 69 genes with an FDR<0.1 in the TADA-mega analysis obtained a simulated p-value of less than 0.006 (median *P*=1×10^−3^), with *CHD8* obtaining a p-value of 9×10^−7^ (**Fig. 4b**).

**Fig. 4.**
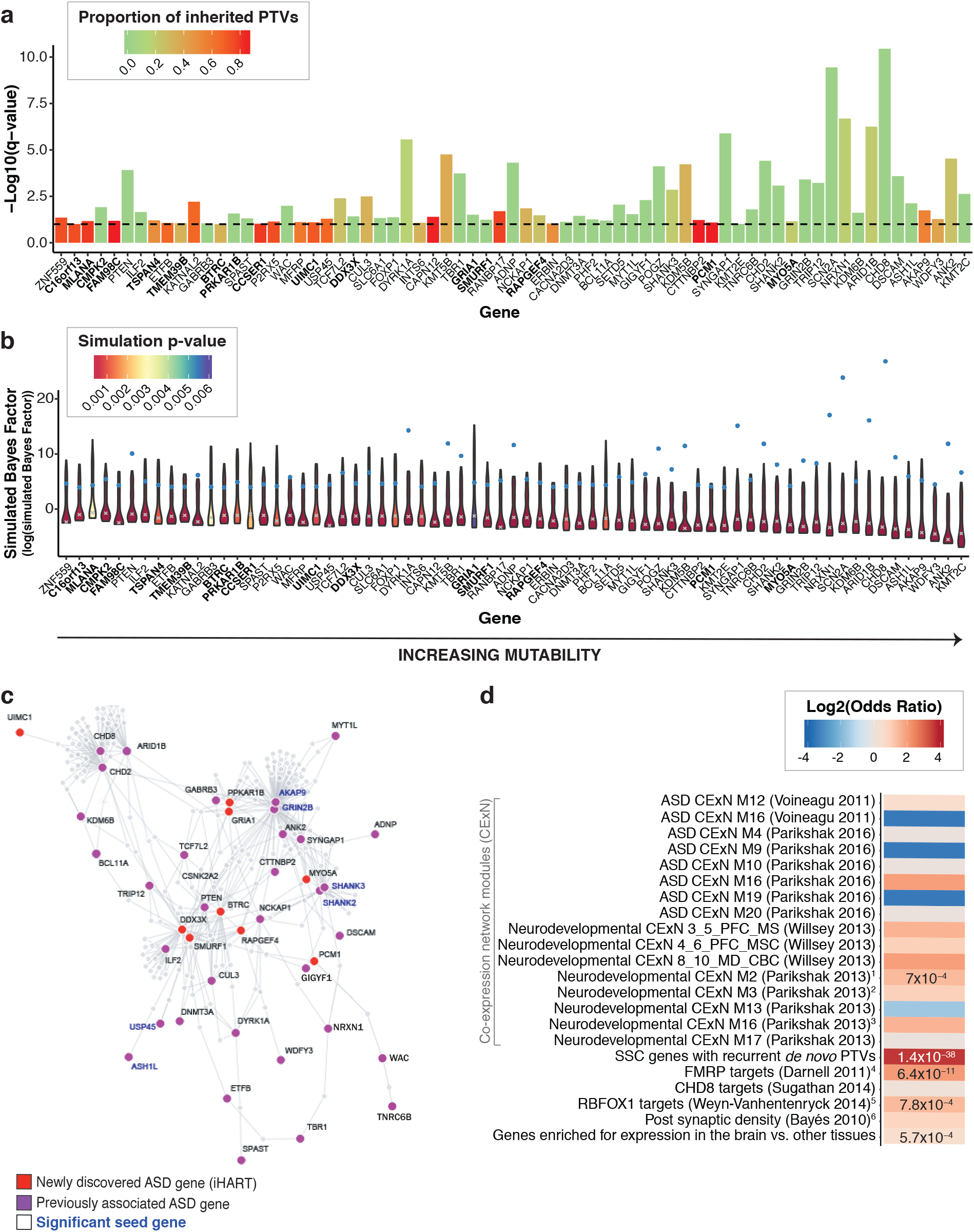
69 ASD-risk genes identified by the TADA mega-analysis. **(a)** The 69 genes with an FDR<0.1 are displayed in order of increasing mutability and the 16 novel genes identified in the iHART mega-analysis are in bold. The per-gene TADA FDR is displayed as a bar reaching the - log10(q-value), so higher bars have a lower FDR; the dashed horizontal line marks the FDR=0.1 threshold; and the proportion of inherited PTVs for each gene (inherited PTVs/(inherited PTVs + *de novo* PTVs + *de novo* missense-3 + *de novo* small deletions)) is displayed. **(b)** The 69 genes with an FDR<0.1 are displayed in order of increasing mutability and the 16 novel genes identified in the iHART mega-analysis are in bold. Violin plots of the simulated Bayes Factors (displayed as log(simulated Bayes Factor), 111 quantiles from the 1.1 million simulations) for each gene. The grey x indicates the median of the simulated Bayes Factors for that gene and the blue dot is the Bayes Factor obtained in the iHART TADA-mega analysis. The violin plots are filled according to their simulation p-value (max p-value=0.006). Genes with a very large distance between the median simulated Bayes Factor and the observed TADA-mega analysis Bayes Factor (e.g., CHD8) are highly secure genes with a very low probability of having achieved the observed Bayes Factor by chance alone given the input family structures. **(c)** The indirect PPI network formed by the 69 ASD-risk genes identified by TADA with an FDR<0.1 with proteins encoded by previously known ASD-risk genes (by the TADA mega analysis in Sanders *et al*. (*22*)) shown in purple and proteins encoded by newly identified ASD-risk genes in the iHART TADA mega analysis shown in red. This network includes 9 of the 16 novel ASD-risk genes. The gene labels for the six significant seed genes are bold and blue. The resulting indirect PPI was significant for two connectivity metrics – seed indirect degrees mean *P*=0.016 and the CI degrees mean *P*=0.024. **(d)** Gene-set enrichment results for the 69 ASD-risk genes displayed by the log2(odds ratio), with p-values listed for gene sets surviving multiple test correction (Bonferroni correction for the 22 gene sets tested or *P*<0.002); the SSC DN PTV recurrent gene set was included as a positive control. In addition to the gene set “genes enriched for expression in the brain vs. other tissues” which contains almost all of the 16 novel ASD-risk genes, six additional gene-sets contain one or more of the 16 novel ASD-risk genes: (1) *TMEM39B* and *PCM1*, (2) *CCSER1* and *UIMC1*, (3) *BTRC, PRKAR1B*, and *MYO5A*, (4) *RAPGEF4* and *MYO5A*, (5) *BTRC*, (6) *DDX3X, GRIA1, RAPGEF4*, and *MYO5A*.

**Table 1.**
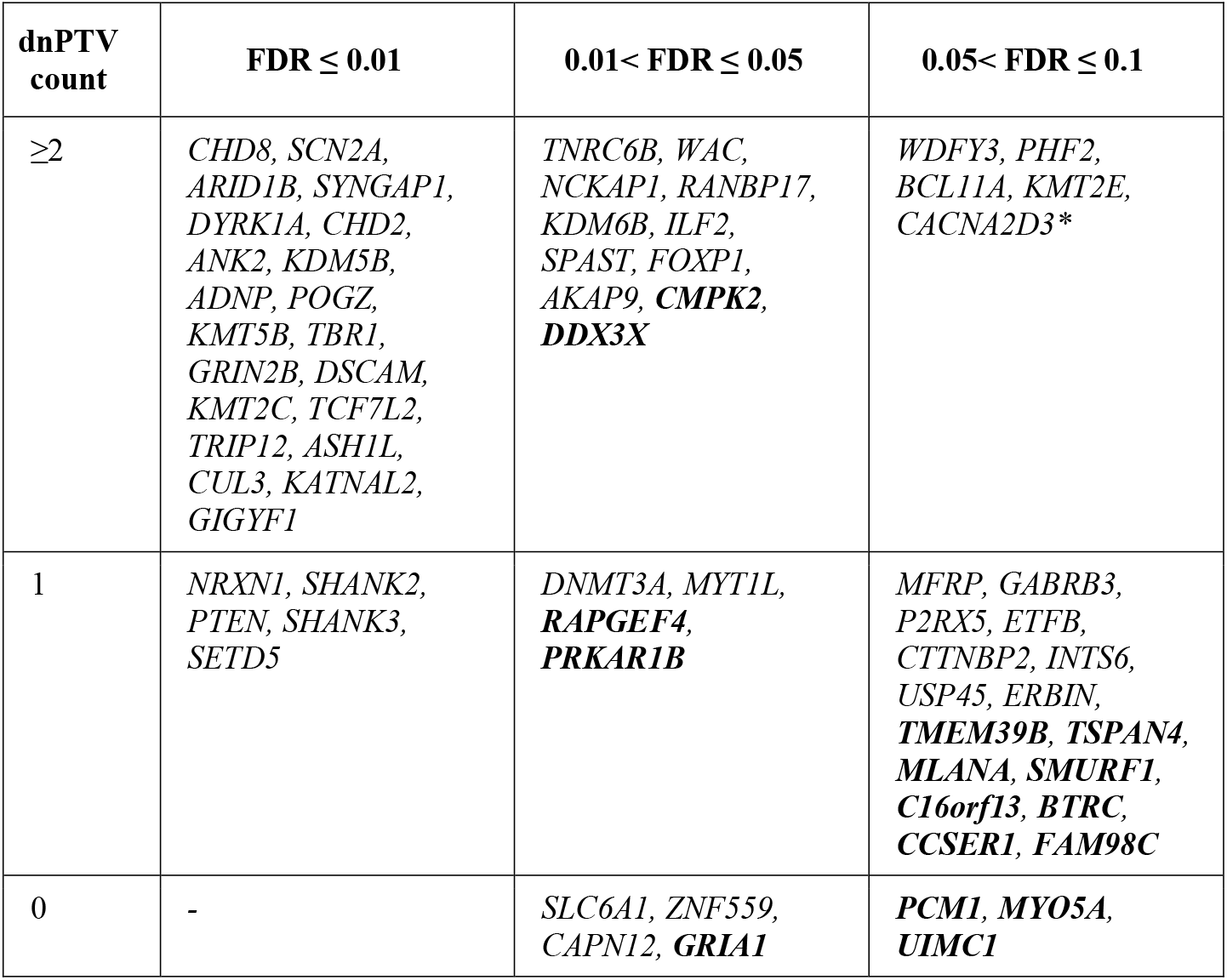
69 ASD-risk genes, including 16 novel genes, identified in the iHART TADA mega-analysis. All 69 genes significantly associated with ASD-risk (with an FDR < 0.1) by the iHART TADA mega-analysis are displayed by the number of *de novo* PTVs identified in the gene. The 16 newly ASD-associated genes are in bold. *The *CACNA2D3* gene was significantly associated with ASD in the iHART mega-analysis but not the previous TADA mega-analysis(22); however, it was previously reported in De Rubeis *et al.(18)* and thus is not considered as a novel ASD-risk gene.

Given the lower relative risk estimated for inherited PTVs(*18*), we relaxed the statistical threshold to an FDR <0.2 (n=119), confirming 84 genes previously identified at this threshold(22). Additionally, we identify 35 genes that had not reached this threshold (FDR<0.2) in the previous study(*22*), 15 of which had the majority (≥70%) of their qualifying risk variants represented by inherited PTVs compared to only 8 of the 84 genes previously identified at this threshold (Fisher’s Exact Test, *P*=7.45×10^−5^, OR=6.98, 95% CI: 2.39-21.96). Consistently, for these 35 genes, the current study obtains higher inherited PTV Bayes Factors as compared to those obtained in the previous TADA mega-analysis in largely simplex families(*22*) (Kruskal–Wallis test, *P*=0.0003, **Fig. S11a**). For five of these 35 genes (*PCM1, STARD9, GRM6, RHPN1*, and *SLC10A1*) and two of the remaining 84 genes (*CTTNBP2* and *ZNF559*) that were previously observed with FDR<0.2, the largest association signal is from inherited PTVs. Thus these 35 genes are enriched for genes whose association signal is primarily driven by inherited PTVs (Fisher’s Exact Test, *P*=0.02, 0R=6.70, 95% CI: 1.03-73.81) (**Methods**), further indicating that there is a substantial, previously under-represented signal from rare inherited variants.

Comparison of the iHART TADA-mega analysis to the previously published findings(*22*) identified 16 newly significant (FDR<0.1) ASD risk genes plus *CACNA2D3* which was previously reported as an ASD risk gene(*18*) (**Table 1, Fig. S10**). However, we also failed to replicate 13 of the genes previously published with an FDR<0.1(*22*) (**Fig. S10**). The q-values for these 13 genes were borderline significant in iHART (**Fig. S10a**), and their simulation p-values were greater (min p-value=0.01, max p-value=0.06, **Fig. S10c**) than those of the securely implicated 69 ASD risk genes, including the 16 newly significant genes (min p-value=0.001, max p-value=0.006) (**Fig. S10d, Fig. 4b**).

### Biological insights from known and novel ASD genes

Gene-set enrichment (**Methods**) of the 69 high-confidence ASD risk genes identified enrichment in a highly co-expressed group of transcriptionally co-regulated genes active during human cerebral cortical neurogenesis (Module 2)(*42*), FMRP targets(*58*), RBFOX1 targets(*44*), and genes enriched for expression in the brain vs. other tissues (**Methods; Fig. 4d**). Analysis of single cell sequencing data from human brain reveals enrichment in mid-gestation and adult glutamatergic projection neurons for both the high confidence and newly identified ASD risk genes (**Methods; Fig. S12**). Many of the 16 new ASD risk genes from this study fall into gene sets of interest. Pathway analysis revealed three biological pathways containing these genes, including negative regulation of synaptic transmission (*RAPGEF4*), learning and memory (*GRIA1* and *PRKAR1B*) and organelle organization (*PCM1* and *MYO5A*) (**Fig. S11b**). Other examples include *PRKAR1B* (q-value=0.026), which is in a gene co-expression module (Module 16) comprised of structural synaptic proteins that are highly co-expressed during human cerebral cortical neurogenesis and in which SFARI ASD risk genes are overrepresented(*42*); and three genes that are found in the post synaptic density (PSD) of the human neocortex(*59*): *GRIA1* (q-value=0.031), *RAPGEF4* (q-value=0.033), and *DDX3X*(q-value=0.038). *RAPGEF4* is also a known FMRP target(58) and was previously identified as an ASD candidate gene based on five families with segregating PTVs(*60*). *DDX3X* was recently reported to account for 1%-3% of unexplained intellectual disability in females(*61*). Finally, 9 of these 16 new ASD risk genes form a significant indirect PPI network (Methods; seed indirect degrees mean permutation *P*=0.016, and CI degrees mean *P*=0.024) (**Fig. 4c**).

### Candidate genes harboring high-risk inherited variation form a significant network

We then asked if the proteins encoded by the 98 candidate genes harboring PTVs and SVs transmitted to all affected individuals interact with the 69 ASD-risk genes identified in the TADA mega-analysis (FDR<0.1). The resulting PPI network formed by these 165 unique genes is significant for the direct edges count (*P*=0.036), the seed direct degrees mean (*P*=0.046), and the CI degrees mean (*P*=0.005) (**Fig. 5a, Fig. S11c**). This network reveals interactions between genes with different levels of statistical support, ranging from high-risk inherited candidate genes, established ASD risk genes, and new ASD-risk genes, which provides evidence that at least some of these 98 candidate genes are true ASD risk genes.

**Fig. 5.**
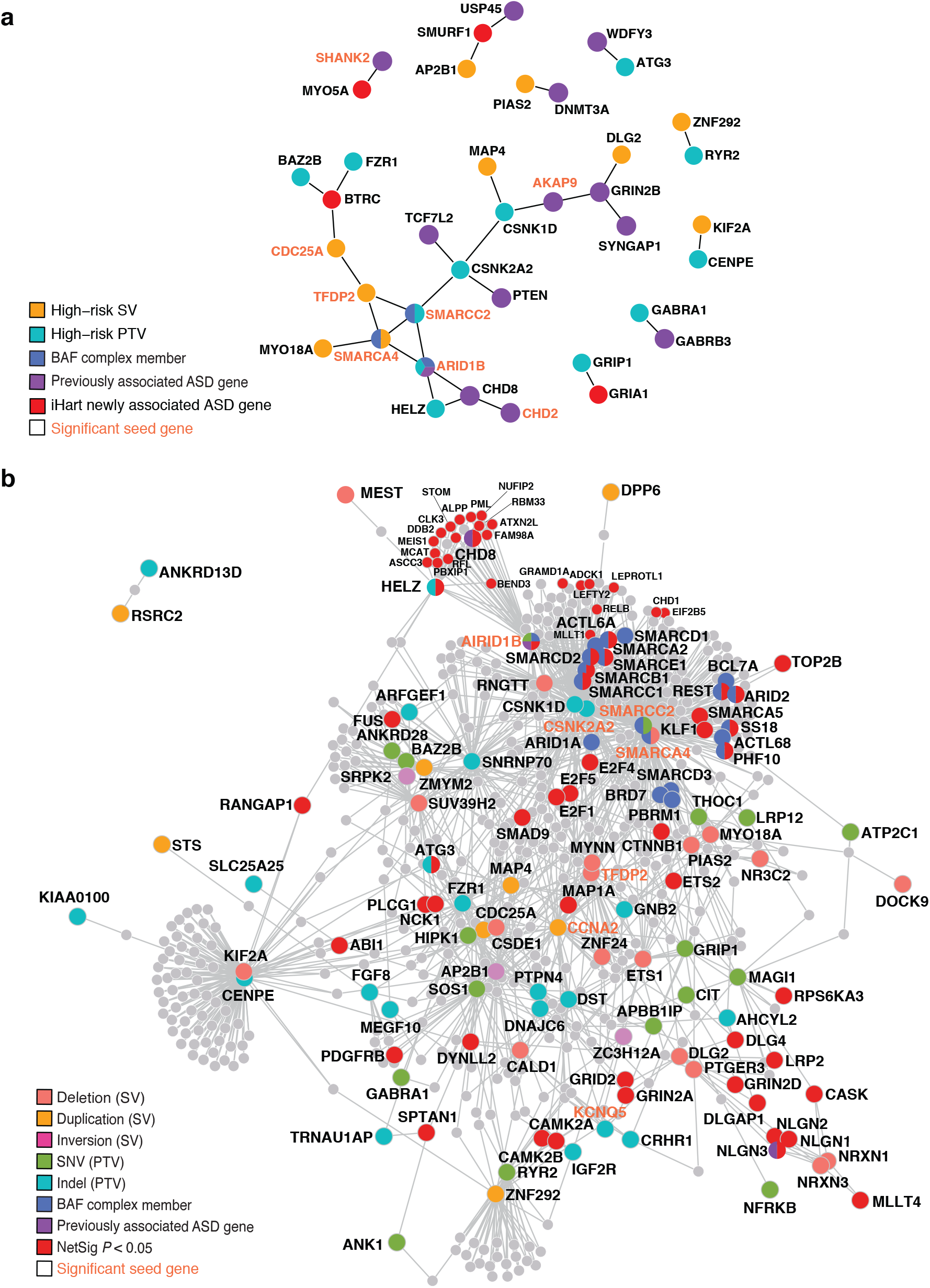
Direct protein-protein interaction network formed by ASD-risk genes. **(a)** The direct protein-protein interaction network (identified by DAPPLE) formed by intolerant genes harboring PTVs or SVs transmitted to all affected (and no unaffected) members of an AGRE family (98 genes) and ASD-risk genes identified in the TADA mega-analysis (69 genes, FDR<0.1). The direct protein-protein interaction network formed by these 165 genes (2 genes in both categories), is significant for three connectivity metrics: the direct edges count (*P*=0.036), the seed direct degrees mean (*P*=0.046), and the CI degrees mean (*P*=0.005). Each encoded protein is colored according to the type of variant class identified; proteins encoded by a gene with a high-risk inherited SV are shown in gold and PTVs are shown in teal. Proteins encoded by an ASD risk gene by the TADA mega analysis in Sanders *et al*. (*22*) are shown in purple, proteins encoded by a newly identified ASD-risk gene by the iHART TADA mega analysis are shown in red, proteins belonging to the BAF complex are shown in blue, and any protein falling in more than one category is colored with all categorical colors that apply (e.g., ARID1B). The gene labels for significant seed genes are bold and orange. **(b)** The indirect protein-protein interaction networks (identified by DAPPLE) formed by intolerant genes harboring PTVs or damaging SVs (promoter or exon disrupting) transmitted to all affected (and no unaffected) members of an AGRE family. Each encoded protein is colored according to the type of variant class identified. Proteins encoded by an ASD risk gene identified in the TADA mega analysis in Sanders *et al*. (*22*) are shown in purple. For visualization, protein symbols for some NetSig significant genes (*P* < 0.05) are displayed in a smaller font in the upper right hand corner of the network. Any protein falling in more than one category is colored with all categorical colors that apply. The gene labels for significant seed genes are bold and orange.

Given that a large number of predicted ASD-risk genes remain unidentified(*24*), we next applied NetSig(*62*), a newly developed method that identifies high probability candidate genes by integrating PPI and association statistics. We identified 596 genes that were significantly directly connected to ASD-risk genes (**Methods; Table S12**), 38 of which are enriched in a developmental co-expression module previously shown to be enriched for ASD-risk genes harboring *de novo* variants (module M2(*42*), *P* = 0.0003, OR = 1.98; 95% confidence interval = 1.37-2.81). Proteins in both the direct and indirect PPI networks seeded by high-risk inherited PTVs directly bind ASD-risk genes more than expected by chance (*P* = 0.02; OR=12.80; 95% confidence interval = 1.07-111.92) and are also enriched in indirect PPI networks (*P* = 4.24×10^−16^; OR=4.90; 95% confidence interval = 3.45-6.85) (**Fig. 5b; Methods; Extended results**).

## Discussion

To date, *de novo* variants have provided compelling evidence for dozens of ASD-risk genes. Here we used WGS to identify more than a dozen new genes that are significantly associated with ASD-risk, the majority of which are due to a contribution from rare inherited mutations. The identification of 16 novel ASD-risk genes was facilitated by exploiting a cohort ascertained for families containing two or more children with ASD where inherited risk variants are likely to contribute to the observed ASD recurrence(*10, 24, 29*). We also identified genes (pLI≥0.9) harboring inherited damaging variants transmitted to all affected children and not transmitted to any unaffected children that form a PPI network. This PPI network is seeded by known ASD-risk genes and members of the BAF complex, and also enriched for proteins that interact with additional ASD-risk genes, many of which are involved in cortical neurogenesis(*42*). This is supported by the single cell sequencing data which reveals expression of many of these ASD risk genes in developing glutamatergic neurons (**Fig. S12**). Further, the observation that the high-confidence novel ASD-risk genes (FDR<0.1) and additional candidate ASD-risk genes (FDR<0.2) are enriched for inherited variation is logically consistent with the fact that previous ASD studies have primarily focused on simplex ASD families, where *de novo* variants are known to have a critical contribution to ASD risk.

We employed WGS to enable the detection of non-coding variants and structural variation at high resolution. We identified one AGRE/iHART family with a likely pathogenic SNV in the *NR3C2* gene and a second family with a structural variant (deletion) disrupting the promoter region for this gene. The shared phenotypic features amongst the variant carriers is consistent with a new syndromic form of ASD (**Table S3**). We were able to infer biological importance of this *NR3C2* putative regulatory deletion given its open chromatin state in human developing brain (ATAC-seq(*37*)) and phenotypic concordance to the family harboring the coding PTV. Otherwise, more broadly, we found no global enrichment for non-coding variation in promoters – structural variant or otherwise – in affected vs. unaffected children. Consistently, a previously published genome-wide investigation of 53 simplex families found a small enrichment (*P*=0.03) for private and DN disruptive variants in fetal brain DNase I hypersensitive sites in probands. However, this signal was limited to DNase I hypersensitive sites within 50Kb of genes that have been previously associated with ASD-risk(48). Advances in methods for analysis of the non-coding genome, similar to what has been done to identify functional PTVs (e.g., constraint metrics such as pLI), are necessary to improve power for identifying non-coding risk variants.

As previous studies have shown(*27*), inherited variation alone does not explain all instances of ASD within multiplex families. Despite finding no global excess of damaging RDNVs in ASD cases in the study, we do identify PTV and Mis3 RDNVs in previously established ASD risk genes, including: *CHD8, SHANK3*, and *PTEN* (**Table S10**). Given our success in uncovering many ASD risk genes whose signal is derived at least partially from inherited variation, it is clear that even modest increases in sample sizes from families with multiple affected children will confirm many new genes. Our machine learning classifier, Artifact Removal by Classifier (ARC), will also allow increases in sample sizes when only LCL-derived DNA is available by distinguishing next-generation sequencing and cell line artifacts from true *de novo* variation. As sample sizes grow, we can confirm whether our observed differences between simplex vs. multiplex families are generalizable, but our data suggest substantial differences in their genetic architecture. Furthermore, with larger cohorts, we may be able to classify risk genes based on inheritance – (1) *de novo* (2) inherited or (3) *de novo* + inherited – to establish if these distinct gene classes are associated with phenotypic severity or specific biological pathways. The iHART portal (http://www.ihart.org/home) provides researchers access to these data, facilitating additional analyses of these samples and integration with future cohorts.

## Supplementary Materials

Materials and Methods

Figures S1-S13

Tables S1-S16

## Acknowledgements

We thank Stephanie N. Kravitz, Cheyenne L. Schloffman, Min Sun, Tor Solli-Nowlan, Virpi Leppa, Hyejung Won, Sasha Sharma, Marlena Duda, Greg Madden McInnes, and Ravina Jain for additional technical support for this research and we thank the New York Genome Center for conducting sequencing and initial quality control.

## Author contributions

E.K.R and L.P.C designed the analytical plans, implemented the analysis pipelines, and interpreted results. J.K.L was responsible for selecting and submitting samples for sequencing. E.K.R, J.J, L.W, and J.K.L performed quality control checks. L.W wrote scripts for data processing and helped interpret results. L.P.C, D.K, J.J, and E.K.R developed ARC. C.H interpreted results and ran TADA-simulations. J.J. and D.P.W designed the access systems. J.J performed joint genotyping, VCF annotation, and data transfers. L.P.C and D.K processed SVs and L.P.C wrote the SV cross-algorithm comparison pipeline. D.P.W initially conceived of the idea for the study and identified funding. E.K.R and D.H.G took the lead in writing the manuscript, and all authors reviewed and approved the manuscript. D.H.G and D.P.W supervised the experimental design and analysis and interpreted results.

## Data and materials availability

The whole-genome sequencing data generated during this study are available from the Hartwell Foundation’s Autism Research and Technology Initiative (iHART) following request and approval of the data use agreement available at http://www.ihart.org/home. Details about the format of the data, access options, and access instructions are included in the Supplemental material of this manuscript.

